# Phylodynamic model adequacy using posterior predictive simulations

**DOI:** 10.1101/255596

**Authors:** Sebastian Duchene, Remco Bouckaert, David A. Duchene, Tanja Stadler, Alexei J. Drummond

## Abstract

Rapidly evolving pathogens, such as viruses and bacteria, accumulate genetic change at a similar timescale over which their epidemiological processes occur, such that it is possible to make inferences about their infectious spread using phylogenetic time-trees. For this purpose it is necessary to choose a phylodynamic model. However, the resulting inferences are contingent on whether the model adequately describes key features of the data. Model adequacy methods allow formal rejection of a model if it cannot generate the main features of the data. We present TreeModelAdequacy (TMA), a package for the popular BEAST2 software, that allows assessing the adequacy of phylodynamic models. We illustrate its utility by analysing phylogenetic trees from two viral outbreaks of Ebola and H_1_N_1_ influenza. The main features of the Ebola data were adequately described by the coalescent exponential-growth model, whereas the H_1_N_1_ influenza data was best described by the birth-death SIR model.

---

Phylogenetic trees depict the evolutionary relationships between groups of organisms. In the context of infectious diseases, pathogen genetic data can be used to infer such trees. By assuming a substitution model and including independent information about time, one can calibrate the molecular clock to obtain time-trees, where the branch lengths correspond to units of time, and internal nodes of the tree represent the timing of divergence events. For rapidly evolving viruses and bacteria it is possible to use the sampling times as time-calibrations (Drummond et al. 2003; Rieux and Balloux 2016). In these organisms, genetic change and epidemiological or ecological processes occur over a similar timescale. Thus, time-trees can be informative about epidemiological dynamics, a field of research known as phylodynamics (Holmes et al. 1993; Grenfell et al. 2004; Kühnert et al. 2011; Volz et al. 2013).

Phylodynamic models describe the distribution of node times, branch lengths, and sampling times. In full Bayesian phylogenetic analyses, the tree, parameters for the molecular clock, the substitution model and the phylodynamic model can be estimated simultaneously using molecular data. In this Bayesian framework, the phylodynamic model is effectively a ‘tree prior’ (e.g. du Plessis and Stadler 2015). The simplest phylodynamic models are the coalescent exponential-growth (CE) and the coalescent constant-size (CC). These two models have very different expectations about the shape of phylogenetic trees, with exponentially growing populations tending to produce trees with longer external branches than those evolving under constant population sizes (O’Meara, 2012; Volz et al., 2013). Alternative phylodynamic models are the birth-death models, which include a parameter to describe the sampling rate, so they have an expectation on the number of taxa and their distribution over time (Stadler 2010; Stadler et al. 2012).

Choosing an appropriate model is important to draw reliable inferences from parameters of interest. For instance, the CE and the constant birth-death (BD) models can estimate the basic reproductive number, *R_0_* (Frost & Volz, 2010; Stadler et al., 2012; Volz et al., 2013), which is defined as the average number of new cases that a single case will generate over the course of its infection in a fully susceptible population (Anderson and May 1979, 1992). Failing to account for complex epidemiological dynamics can bias the estimate of this key parameter (Stadler et al. 2014; Alkhamis et al. 2016; Ratmann et al. 2016). A Bayesian approach to selecting a phylodynamic model is to estimate marginal likelihoods for a pool of models and selecting that with the highest marginal likelihood (Baele et al. 2012; Baele et al. 2016), but it is also possible to obtain weighted averages of parameter estimates based on the support for each model (e.g. Baele, Li, Drummond, Suchard, & Lemey, 2013; R. Bouckaert & Drummond, 2017; Huelsenbeck, Larget, & Alfaro, 2004; Li & Drummond, 2012).

## Model adequacy in phylogenetics

Model selection methods only allow a relative comparison of a set of models, but they cannot determine whether any of the models in question could have generated key features of the data at hand (i.e. absolute model fit). Such information however is key to avoid unreliable inferences from a model and to improve our understanding of the biological processes that produced the data. Absolute model fit can be assessed via model adequacy methods, where a model is considered ‘adequate’ if it is capable of generating the main features of the empirical data. Consequently, model adequacy allows the user to formally reject a model or to identify aspects of the data that are poorly described, instead of ranking it with respect to other models, as is the case with model testing (e.g. Goldman 1993; Bollback 2002; Ripplinger and Sullivan 2010; Brown 2014).

Model adequacy is typically conducted by fitting a model to the empirical data, and generating synthetic data from the model in question, a procedure that is similar to a parametric bootstrap (Goldman 1993). The adequacy of the model is determined depending on whether the synthetic data are similar to the empirical data, according to a descriptive test statistic (Gelman and Shalizi 2013; Gelman et al. 2014). The test statistics should summarise key aspects of the data or a combination of the data and parameter estimates (Gelman et al. 1996). Examples of test statistics that have been used to assess the substitution model include the multinomial likelihood or a measure of compositional homogeneity (Goldman 1993; Huelsenbeck et al. 2001; Foster 2004). The joint clock model and tree prior key can be assessed using the expected number of substitutions in individual branch lengths of the tree as test statistics (Duchêne et al. 2015). For DNA barcoding the number of OTUs and multinomial likelihood have been shown to be effective test statistics (Barley and Thomson 2016). Phylodynamic and diversification models are fitted to phylogenetic trees and their parameters depend on the distribution of nodes, such that some useful test statistics include the ratio of external to internal branch lengths, the tree height, and measures of phylogenetic tree imbalance (Revell et al. 2005, 2008; Drummond and Suchard 2008; Höhna et al. 2015). Clearly, designing test statistics is not trivial, but they should attempt to explicitly test some of the assumptions of the model. For example, the CE and CC models have different expectations of the ratio of external to internal branch lengths, such that this may be a useful test statistic.

## Bayesian model adequacy

Bayesian model adequacy consists of a posterior predictive framework (Rubin 1981, 1984; Bollback 2002; Brown 2014a, 2014b; Lewis et al. 2014; Höhna et al. 2017). The posterior distribution of the model in question is approximated given the empirical data, for example using Markov chain Monte Carlo (MCMC). Samples from the MCMC are drawn to simulate data sets under the model used for the empirical analysis. For example, the posterior distribution of the growth rate and population size parameters of the CE model can be sampled to simulate phylogenetic trees. Such simulations are known as posterior predictive simulations. Test statistics are then calculated for every posterior predictive simulation (i.e. for every simulated tree) to generate a distribution of values according to the model. A posterior predictive probability, similar to the frequentist *p*-value, can be calculated by determining where the value of the test statistic for the empirical data (i.e. the empirical phylogenetic tree) falls with respect to the posterior predictive distribution (Gelman et al. 2014). Following Gelman et al. (2014) we refer to the posterior predictive probability as *p_B_* to differentiate it from the frequentist *p*-value. A useful guideline to determine whether the model is adequate is to determine whether a test statistic within the 95% credible interval (Bollback 2002; Brown 2014a). This approach is sometimes conservative, particularly when test statistics do not follow a Gaussian distribution, and other methods of calculating posterior predictive probabilities are also possible (Gelman et al. 2014; Höhna et al. 2017). Combining multiple test statistics leads to multiple testing, which can be addressed by using a multivariate *p_B_* values (Drummond and Suchard 2008). However, Gelman et al. (2014) suggest considering each test statistically separately to assess individual aspects of the model and the data, which is the approach taken here.

Bayesian model adequacy is similar to Approximate Bayesian Computation (ABC) techniques in that both methods use test statistics from simulated data. The aim of ABC is to approximate the posterior by comparing test statistics from simulations from the prior and the empirical data (Csilléry et al. 2010; Ratmann et al. 2012; Poon 2015), which sometimes leads to biases for model testing (Robert et al. 2011). In contrast, in model adequacy the simulations are generated from the posterior distribution and they are not used to approximate the posterior. In spite of these differences, test statistics developed for ABC can be useful to assess model adequacy.

## “TreeModelAdequacy” package in BEAST2

We implemented a computational framework to assess the adequacy of phylodynamic models as a package for BEAST2 (Bouckaert et al., 2014). Analyses as outlined in Fig 1 are easy to set up through BEAUti, the graphical user interface for BEAST, to generate an xml file with the tree, the model, and test statistics. Our package, TreeModelAdequacy (TMA), takes a tree with branch lengths proportional to time. The tree can be a summary tree from BEAST2, or estimated using a different method.

**Fig 1.**
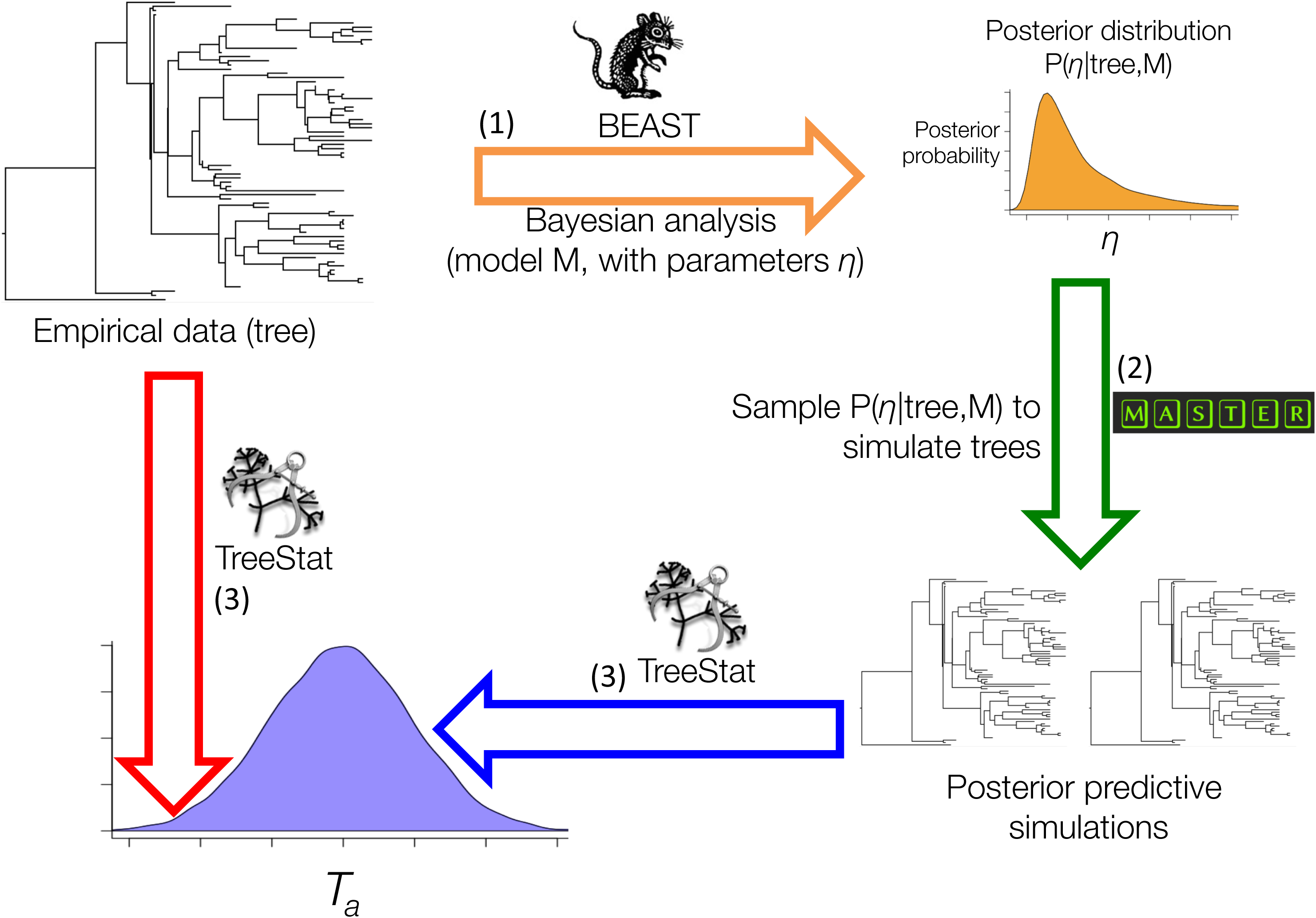
Posterior predictive simulation framework implemented in the TreeModelAdequacy package. Step (1) consists in Bayesian analysis in BEAST under model M to estimate the posterior distribution of parameters *η*, shown with the arrow and posterior density in orange. In step (2) samples from the posterior are drawn to simulate phylogenetic trees, known as posterior predictive simulations, using MASTER as shown by the green arrow. In step (3) the posterior predictive simulations are analysed in TreeStat to generate the posterior predictive distribution of test statistic *T_a_*, shown by the blue arrow and probability density. Finally, *T_a_* is also computed for the tree from the empirical data using TreeStat, shown by the red arrow, to calculate a posterior predictive probability (*p_B_*). Test statistics and *p_B_* values can also be computed for trees generated in other programs using TreeModelAdequacyAnalyser, given that the tree from the empirical data and the posterior predictive simulations are provided.

We fit phylodynamic models available in BEAST2 by approximating the posterior distribution of the parameters of the model using MCMC. To generate the posterior predictive simulations, we draw random samples from the posterior for the parameters from the MCMC after removing the burn-in phase, and we simulate phylogenetic trees using stochastic simulations and master equations using MASTER (Vaughan and Drummond 2013) or the coalescent simulator in BEAST2. The last step consists in calculating test statistics for the empirical data and for the posterior predictive simulations, which depends on the TreeStat2 package (available at http://github.com/alexeid/TreeStat2). The user can select a large number of test statistics (Supplementary material). At the end of the analysis, *p_B_* values and quantiles for the posterior predictive distribution are shown, but they can also be visualised in Tracer (available at: http://beast.bio.ed.ac.uk/tracer) or using an R script included in the package. At present, the range of phylodynamic models that can be assessed includes the CC and CE coalescent models, the constant BD with serial sampling (Stadler et al. 2012), and the birth-death susceptible-infected-recovered model (BDSIR; Kühnert et al. 2014).

Our implementation allows parallelisation of the tree simulation step, which can increase computational speed when the simulation conditions require extensive calculations. This step can also be conducted independently on a computer cluster. Our standalone application TreeModelAdequacyAnalyser can also compute test statistics and *p_B_* values, even for trees generated in a different program than BEAST2, given that the posterior predictive trees are provided. TMA is open source and freely available under a LGPL licence. It can be downloaded from BEAUTi2 (part of BEAST2), and the documentation and example files are available at: http://github.com/sebastianduchene/tree_model_adequacy.

To verify our implementation, we conducted a simple simulation experiment. We simulated 100 trees under each of the four phylodynamic models (CC, CE, BD, and BDSIR) using BEAST2 and MASTER and analysed them with the matching model. We assessed their adequacy according to nine test statistics (Supplementary material). The parameters for our simulations were based on analyses of 72 whole genome sequences of Ebola virus (Gire et al. 2014). We found that the *p_B_* values for all test statistics were between 0.025 and 0.975 for about 95% of each set of simulations, indicating that our implementation is correct (Supplementary material).

## Phylodynamic model adequacy in empirical virus data

### West African Ebola virus

We obtained a phylogenetic tree inferred in a previous study from 72 Ebola virus whole genome samples collected during the 2013–2016 epidemic (Gire et al. 2014). The samples were collected from May to July 2014 in Sierra Leone. These data have been used in previous studies to estimate epidemiological parameters, with estimates of *R_0_* ranging from 1.5 to 2.5, depending on the phylodynamic model (Stadler et al., 2014; Volz & Pond, 2014). An important consideration about our analysis is that we assume that the tree topology and divergence time estimates are sufficiently accurate and that the data are informative.

We inferred phylodynamic parameters for the Ebola virus tree using four models; CC, CE, the BD, and the BDSIR (Kühnert et al. 2014). The CE, BD, and BDSIR models can estimate *R_0_* if information about the sampling process (for the birth-death models) or present number of infected individuals (for the coalescent) is available. In this case, we assumed that the sampling proportion was 0.7 (Gire et al. 2014) by fixing this parameter in the BD and BDSIR models. For the remaining parameters of these two models, we used the same prior distributions as in a previous analysis of these data (Stadler et al. 2014). For the CE model we used a Laplace distribution and a 1/*x* distribution as priors for the growth rate and for the effective population size, respectively. We ran an MCMC of 10^7^ steps and generated 1,000 posterior predictive simulations, and we computed four test statistics.

To compare the different models, we calculated *p_B_* for four test statistics; the tree height, the slope ratio of a lineages-through-time (LTT) plot, the ratio of external to internal branch lengths, and the Colless index of phylogenetic imbalance (Fig. 2; Supplementary material). Importantly, the slope ratio of the LTT plot has been found to be informative for inferring epidemiological parameters using ABC (Saulnier et al. 2017).

**Fig 2.**
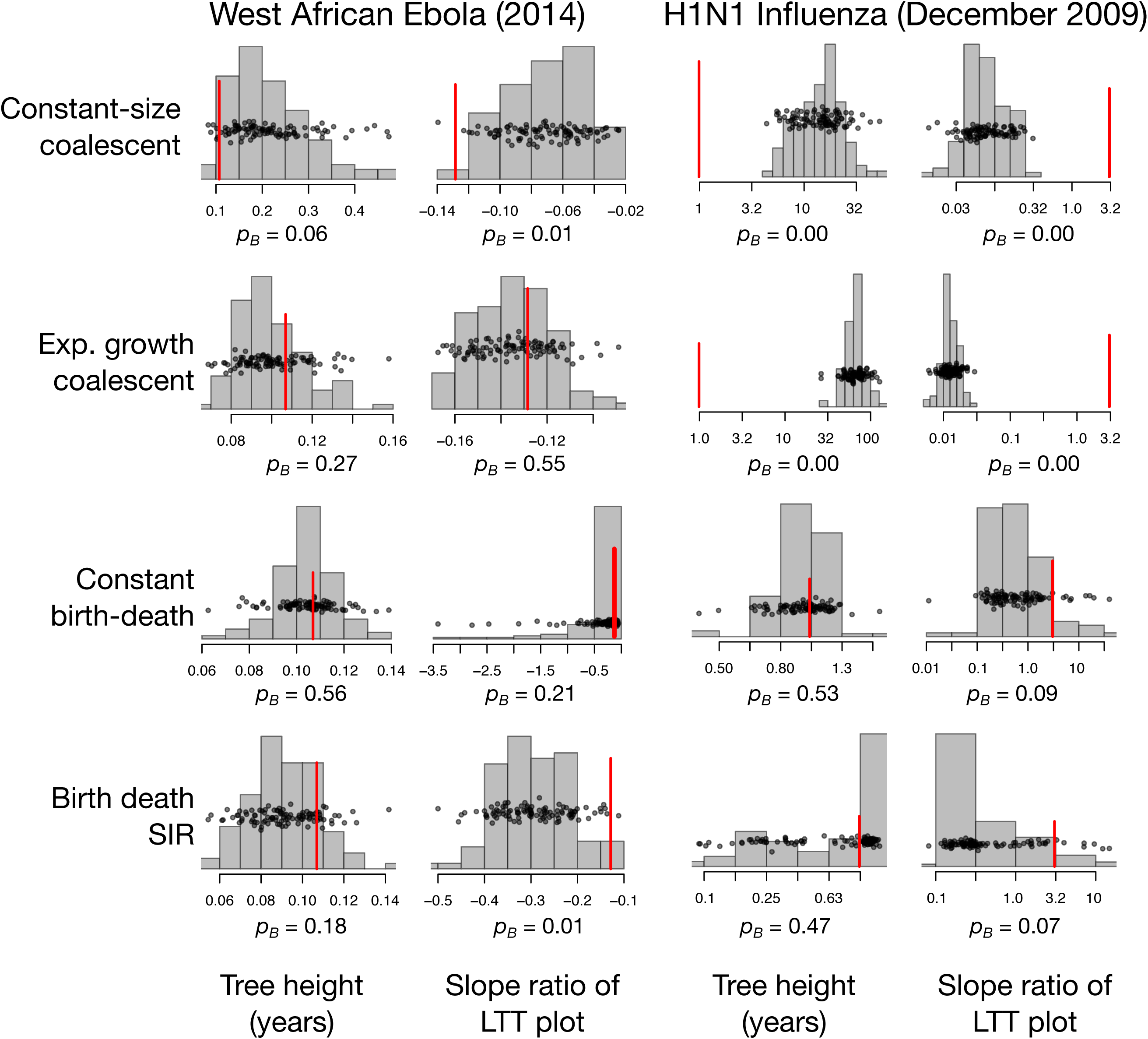
Model adequacy results for the two empirical data sets, West African Ebola and 2009 H_1_N_1_ influenza. The histograms show the distribution of two test statistics, the tree height and the slope ratio of the lineages-through-time (LTT) plot, for the posterior predictive simulations. The black points also show the distribution of the test statistics, and they have been jittered along the *y*-axis to improve visualisation. The red lines denote the value for the tree estimated from the empirical data. The posterior predictive *p*-value, *p_B_*, is shown for each test statistic. A feature of the empirical phylogenetic tree described by a test statistic, such as the tree height, is considered adequately described by the model if the empirical value falls within the 95% quantile range of the posterior predictive distribution, such that the *p_B_* > 0.05. Two more test statistics were computed, the ratio of external to internal branch lengths and the Colless index, which are shown in Supplementary material Fig. S1. Note that for the H_1_N_1_ influenza analyses, the values are shown in a log_10_ scale.

In the CC model the *p_B_* was < 0.05 for all test statistics, with the exception of the tree height at 0.06 (Fig. 2 and Supplementary material Fig. S1). The CE and BD models described these data better, with most *p_B_* values between 0.11 and 0.56. The *p_p_* value for the Colless index in the CE model was the lowest for both of these models, at 0.04. The BDSIR model had overall low *p_B_* values, from to 0.01 to 0.18, with the lowest values found for the ratio of external to internal branch lengths (0.03) and for the slope ratio of the LTT plot (0.01). The fact that most *p_B_* values for the CE and BD models frequently fall near the centre of the posterior predictive distributions is consistent with the rapid spread that Ebola virus was undergoing at the time the sequences were collected.

For the CE, BD, and BDISIR models we can estimate *R_0_* and the infectious period, 1/*δ*. The *R_0_* median estimates were: 1.6 (95% credible interval (CI): 1.12 – 2.2) for the BD, 1.21 (95% CI: 1.1 – 1.5), for the CE, and 1.59 (95% CI: 1.22–1.94) for the BDSIR. Estimates for the infectious period in calendar days were: 5.46 (95% CI: 4.14 – 7.24) for the BD, 2.90 (95% CI: 1.75 – 5.50) for the CE, and 5.21 (95% CI: 4.24–7.02) for the BDSIR. The estimates from these models were very similar and overlapped with those from previous studies (Stadler et al. 2014). Although the BDSIR did not capture some of the main features of these data, this model is similar to the BD when the number of susceptible individuals is very large, which probably explains the overlap in *R_0_* estimates between these models.

### 2009 H_1_N_1_ Influenza

We obtained a phylogenetic tree from a previous study (Hedge et al. 2013), which was estimated from 328 whole genome samples from the 2009 H_1_N_1_ Influenza pandemic. The samples were collected from April to December 2009, such that they encompass a large portion of the duration of the pandemic. We used a similar method as for the Ebola virus phylogenetic tree to fit the four phylodynamic models (CC, CE, BD, and BDSIR). However, instead of fixing the sampling proportion we used an informative prior distribution of the infectious period via the *becomeUnifectiousRate* parameter, with a normal distribution of mean 85 and standard deviation of 15 (corresponding to an infectious period of about 4.45 days).

The CC and CE models had *p_B_* values of 0.00 for all four test statistics, such that they did not adequately describe any of these aspects of the tree. The BD model had *p_B_* values of 0.53 and 0.09 for the tree height and the slope ratio of the LTT plot, and of 0.00 for the ratio of external to internal branch lengths (Fig. 2 and Supplementary material Fig. S2). In contrast, the BDSIR model overall described the H_1_N_1_ tree better overall than the other three models, with *p_B_* values of between 0.07 and 0.44. This result is consistent with the sampling time of the data, which includes the start of the pandemic and the decline in the number of infections towards the end of the year. We calculated an *R_0_* of mean 3.01 (95% CI: 2.5–3.7) at the start of the pandemic in January that declined to *R_0_* < 1 around June, when infectious spread was lower. This estimate is similar to those made in previous studies based on census data (Forsberg White et al. 2009), but in the higher range of those based on the CE model for samples collected in early stages of the pandemic (e.g. Hedge et al. 2013). For comparison, the *R_0_* estimate from the BD model, which appeared inadequate, was substantially lower, with a mean of 1.02 (95% CI: 1.00 – 1.03).

## Conclusion

Model adequacy methods are useful to understand the biological processes that generate the data, such as the evolutionary branching process. For example, our approach reveals that in July of 2014 the West African Ebola outbreak was still growing exponentially and that the 2009 H_1_N_1_ influenza virus pandemic had evidence of a depletion of susceptible individuals in December. In some cases, identifying models that indadequately describe key aspects of the data may improve estimates of parameters of interest, such as *R_0_* in our H_1_N_1_ influenza analyses. One consideration of our approach is that it requires an accurate estimate of a single phylogenetic tree. Clearly, the phylogenetic tree should be inferred using informative sequence data and the sensitivity to the prior should be carefully examined (Ritchie et al. 2016; Boskova et al. 2018; Möller et al. 2018), which is also the case for any Bayesian analysis. For example, a tree estimated from uninformative sequence data will be driven by the prior and will necessarily appear to be adequately described by the matching model, potentially leading to increased rates of type 2 errors. A limitation of our method is that it does not account for phylogenetic uncertainty, which can be addressed by comparing sets of trees from the posterior with those from the posterior predictive distribution. However, this approach will require the development of new test statistics and model assessment criteria. Model adequacy in phylogenetics will benefit from further development of methods to assess more sophisticated phylodynamic models, such as those that account for population structure (Kühnert et al. 2016; Müller et al. 2017a, 2017b; Volz and Siveroni 2018), and techniques to improve the interpretation of *p_B_* values for test statistics that are not normally distributed, such at the Colless index. Model adequacy software, such as TMA, will be key to address these questions.

## Acknowledgements

The authors thank Timothy Vaughan for valuable discussions. SD was supported by a McKenzie fellowship and a Dyason grant from the University of Melbourne. AJD was supported by a Rutherford fellowship (http://www.royalsociety.org.nz/programmes/funds/rutherford-discovery/) from the Royal Society of New Zealand. TS was supported in part by the European Research Council under the Seventh Framework Programme of the European Commission (PhyPD: grant agreement number 335529).

## Author contributions

SD, RB, and AJD wrote the computer code. SD analysed the data. TS, SD, and AJD designed the experiments. SD wrote the manuscript with input from all the authors.

## Supplementary figure captions

**Fig S1.** Model adequacy results for Ebola virus data with four test statistics. The colours and symbols match those in Fig 2.

**Fig S2.** Model adequacy results for H_1_N_1_ influenza data with four test statistics. The colours and symbols match those in Fig 2. Note that the values are shown in log_10_ scale.

